# The Binding Landscape of an Essential Nuclear Hormone Receptor in *C. elegans*

**DOI:** 10.64898/2025.12.14.694179

**Authors:** Alexander L. Sinks, Masako Asahina, Keith R. Yamamoto, Jordan D. Ward, Deborah M. Thurtle-Schmidt

**Author notes:** **Corresponding Author:** Deborah M. Thurtle-Schmidt Biology Department 209 Ridge Rd, Box 5000 Davidson, NC 28031.

## Abstract

Transcription factors (TFs) control gene expression by binding to response elements but distinguishing functional binding sites from non-functional ones remains a major challenge in the field of transcriptional regulation. To probe transcription factor binding and function in a multicellular organism we characterized the binding and transcriptional landscape of NHR-25, a highly-conserved and essential transcription factor in *C. elegans*. Using CRISPR/Cas9, we tagged the essential nuclear hormone receptor, *nhr-25*, with GFP and FLAG at its endogenous locus in *C. elegans* and performed ChIP-seq. We found that NHR-25 binds with two distinct modes: proximal promoter binding and distal enhancer-type binding. Motif analysis of the ChIP-seq peaks showed that the NHR-25 binding motif depended on chromatin context and identified additional enriched motifs, suggesting other putative regulatory partners at composite response elements. By combining ChIP-seq with conditional RNAi knockdown, we characterized *nhr-25’s* transcriptome and identified putative direct targets of NHR-25 binding. Through an 11-bp mutation of the summit of an identified NHR-25 occupied region, we demonstrated that a single response element was responsible for regulating the expression of multiple genes. Our findings provide a comprehensive view of an essential transcription factor’s functional binding landscape at native protein levels, highlighting the intricate nature of response element logic and demonstrating the critical role of a single regulatory element in regulating gene expression.

## INTRODUCTION

Correct transcriptional regulation is key to proper cell identity. Transcriptional factors bind to specific DNA sequences and direct precise gene regulatory networks in distinct cell types. However, knowing where a transcription factor binds in the genome is challenging as only a fraction of the total the transcription factor motifs in the genome are occupied at a time (Ren et al. 2000; Harbison et al. 2004). Convoluting predicting transcription factor binding even further, many sites occupied by transcription factors do not have the cognate motif for the bound transcription factor (Inukai et al. 2017; Mahendrawada et al. 2025). Thus, transcription factors must navigate the chromatin environment and interact with other regulatory factors to accurately respond to signals in specific cell types. Analysis of genome-wide transcription factor occupancy describe two modes of transcription factor binding: promoter proximal and distal binding resulting in action at a distance (Chen et al. 2020). However, these different binding modalities make it difficult to accurately predict a transcription factor target gene from its binding location.

The nematode *C. elegans* is a powerful model organism for genetic and developmental biology, but the highly compact nature of its genome has historically led to the underappreciation of distal regulatory elements. More recent genomic explorations, including the systematic mapping of many transcription factors, description of chromatin states, and cataloging of differentially accessible regions have shed light on the existence and importance of these elements in *C. elegans* (Daugherty et al. 2017; Araya et al. 2014; Jänes et al. 2018; Gerstein et al. 2010; Evans et al. 2016). However, these studies focus on a system-wide approach rather than testing functional impact of transcription factor binding through dissection of a single transcription factor important for development. One such transcription factor is *nhr-25*, a highly conserved nuclear hormone receptor that is an essential gene and necessary for proper development, including molting, vulval, hypodermal, and gonadal cell specification (Gissendanner and Sluder 2000; Asahina et al. 2000; Chen et al. 2004; Asahina et al. 2006). Previous studies have mapped its binding using various methods, including multi-copy tagged integrants (Araya et al. 2014; Shao et al. 2013) and DamID (Katsanos and Barkoulas 2022), but these studies do not capture the binding at endogenous expression levels. Since transcription factor function is dependent on dosage (Noviello 2025), including for *nhr-25* (Ward et al. 2014b), altering expression could alter the binding landscape.

In this study, we performed an integrated analysis of transcriptional regulation in *C. elegans* to reveal functional roles for transcription factor binding and function. Using CRISPR/Cas9 gene editing, we created a strain with a GFP-FLAG tagged, functional *nhr-25* gene, and then profiled NHR-25 binding at endogenous levels in L1 larvae. Specifically, we integrated ChIP-seq identified peaks with genomic chromatin classifications to investigate NHR-25 binding modalities. Additionally, we investigated functional consequences of transcription factor action through knockdown and mutation of *nhr-25*, identifying putative direct targets of and characteristic predictors of functional NHR-25 binding. Finally, through a targeted CRISPR deletion of a *cis*-regulatory response element, we demonstrate that a single regulatory element can be responsible for the expression of multiple genes, highlighting the complex regulatory roles of a key developmental and highly conserved transcription factor in *C. elegans*.

## RESULTS

### The binding landscape of endogenously-tagged NHR-25

To characterize transcription factor binding in *C. elegans*, we profiled binding of a highly-conserved nuclear hormone receptor, NHR-25, at L1. We generated an endogenous *nhr-25::GFP::BioTag::AID*::3xFLAG* (hereafter called *nhr-25::GFP::3XFLAG* for simplicity) knock-in in an *eft-3p::TIR1::mRuby* background to produce a C-terminal translational fusion that labels both *nhr-25* isoforms (Fig. 1A). Homozygote knock-ins were superficially wild type with no observable *nhr-25* mutant phenotypes (embryonic lethality, vulval or molting defects) indicating that the tag did not disrupt gene function and that there was negligible auxin-independent degradation phenotypes (Gissendanner and Sluder 2000; Asahina et al. 2000; Chen et al. 2004; Yesbolatova et al. 2020; Schiksnis et al. 2020). Tagging the *nhr-25* locus with both GFP and FLAG allowed for (1) direct observation of NHR-25 expression in worms prepared for ChIP samples and (2) use of a well-validated antibody for ChIP studies. Direct observation of NHR-25 prior to harvesting for ChIP-seq was critical as *nhr-25* expression is highly oscillatory throughout the molting cycles (Gissendanner et al. 2004; Hendriks et al. 2014). Thus, we wanted to ensure that samples were harvested when NHR-25 was expressed at high levels. Additionally, this tagging strategy was similar to that of a previous NHR-25 ChIP-Seq study, which used integration of a fosmid reporter (Shao et al. 2013). *nhr-25::GFP::3xFLAG* and N2 control animals were harvested for ChIP-Seq at mid L1 (5 hours after L1 release at 25°C) when NHR-25 is expressed in the developing gonad – Z1/Z4 and hypodermis (Fig 1B and C). We conducted two biological replicates in which the IPs show strong correlation between the replicates (r=.95) (Supplemental Fig. S1A), confirming experimental reproducibility.

**Figure 1.**
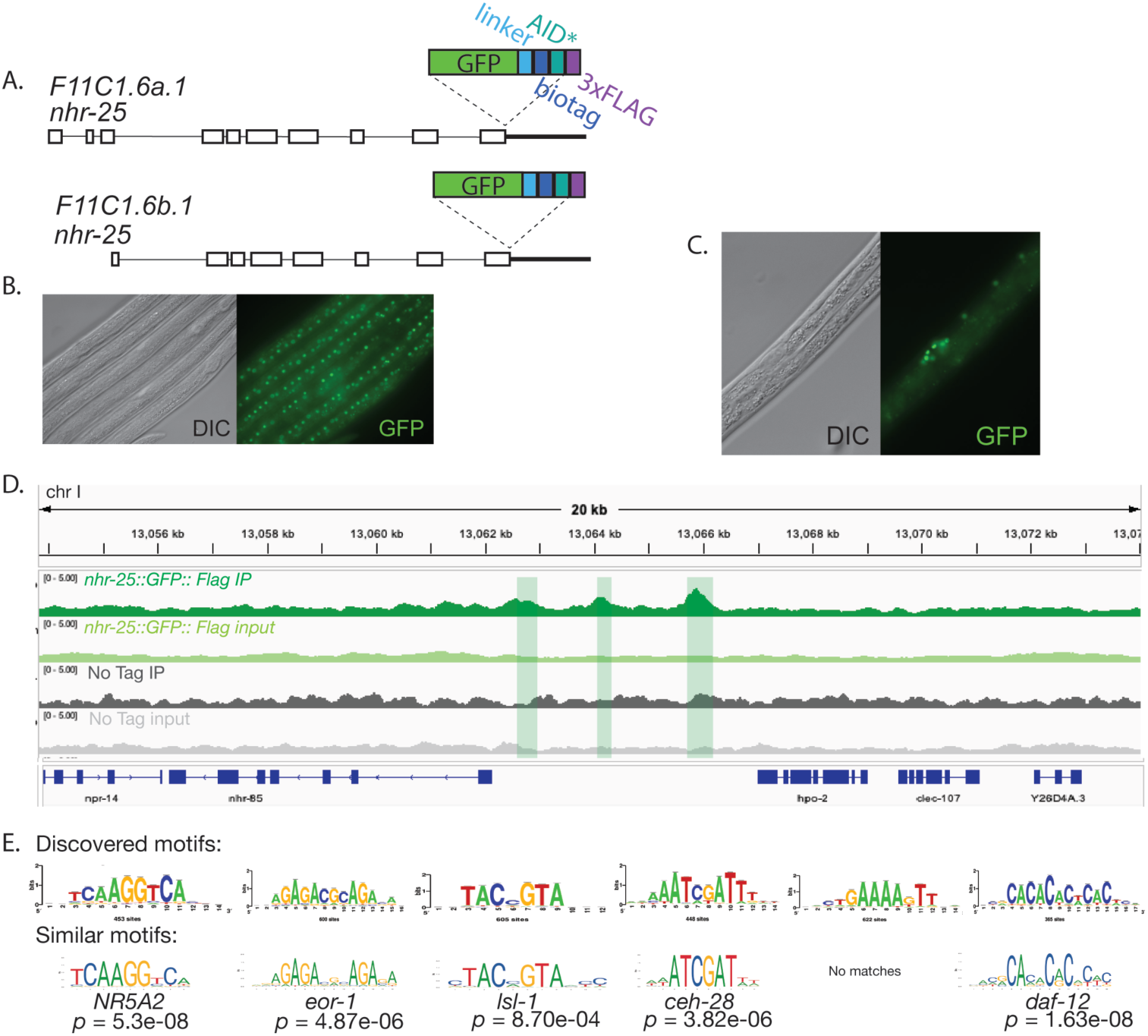
NHR-25 genome occupancy in L1 worms. (A) Schematic of the endogenous *nhr-25* genomic locus following CRISPR insertion of an in-frame *GFP::BioTag::AID*::3xFLAG* shown for both *nhr-25* isoforms. (B and C) Expression of NHR-25 as detected by GFP in the hypodermis (B) and developing somatic gonad (C) in samples used for ChIP-Seq. (D) Representative locus of Chromosome I surrounding *nhr-85*. The top row is the FLAG IP normalized coverage, second row in the input normalized coverage for the FLAG IP, and the bottom two rows show the coverage in the IP and input in a strain where *nhr-25* was not tagged. Screenshots generated with IGV (Thorvaldsdóttir et al. 2013). (E) The top row depicts RSAT *de novo* motif identification and enrichment with highly similar known motifs and respective p-values shown below identified with TOM-TOM (Thomas-Chollier et al. 2012; Bailey et al. 2015).

To generate a genomic binding map for an essential transcription factor at endogenous levels and identify NHR-25 associated genetic loci, we identified sites of enrichment, or peaks, in the *nhr-25::GFP::3xFLAG* IP samples using a stringent filtering process. First, peaks were called in the *nhr-25::GFP::3xFLAG* IP sample using MACS2 with the input sample as the control (Zhang et al. 2008). In addition to the *nhr-25::GFP::3xFLAG* ChIP, we also conducted a ChIP on N2 at L1, in which *nhr-25* was not tagged. Peaks were also called in this “no-tag” dataset, and we disregarded any NHR-25 peaks that overlapped with “no-tag” peaks. Previous studies including work from the modENCODE project have identified regions of the *C. elegans’* genome where a large fraction of transcription factors bind, likely indicating non-specific enrichment (Araya et al. 2014; Teytelman et al. 2013; Gerstein et al. 2010; Moorman et al. 2006). To focus our set on those regions specific to NHR-25, we filtered out any NHR-25 peaks overlapping L1 XOT (Extreme-Occupied Regions) regions which are defined as areas with significant (False Discovery Rate or <1%) transcription factor occupancy across all datasets (Araya et al. 2014). Through this analysis we identified 1140 enriched regions that were specifically identified for the combined NHR-25 ChIP-seq analysis (Table S1). A representative region upstream of *nhr-85*, a nuclear hormone receptor which regulates *lin-4* and is involved in egg-laying pathways (Gissendanner et al. 2004; Kinney et al. 2023), is shown with three enrichment peaks highlighted (Fig. 1D).

Transcription factor binding regions are characterized by several features including open chromatin and enriched sequence-specific motifs particular to that factor. Thus, we sought to validate our NHR-25 L1 endogenous ChIP-seq by investigating the identified peaks. First, we compared the accessibility of the peaks and surrounding areas using L1 ATAC-seq data from a previous study (Jänes et al. 2018). The identified 1140 enriched regions overall showed high chromatin accessibility as would be expected for a transcriptional regulatory region (Supplementary Fig S1B). Additionally, the DNA Binding domain of NHR-25 is highly conserved and binds to the 9bp NR5A2 motif *in vitro* (Gissendanner and Sluder 2000; Chen et al. 2004; Asahina et al. 2006; Ward et al. 2013). Thus, we would expect this motif to be enriched in our identified binding regions. We performed de novo motif discovery and enrichment analysis (Nguyen et al. 2018) of the recovered regions showed a motif highly similar to the NR5A2 motif (p = 5.3e-08), suggesting we have identified a high-confident set of NHR-25 enriched regions at L1 (Fig. 1E). The motif was identified in 40% of enriched regions which is consistent with previously published studies for crosslinked transcription factor occupancy in yeast (Mahendrawada et al. 2025). In addition to the presumptive NHR-25 recognition sequence, five other motifs were identified, once condensed with motif clustering (Castro-Mondragon et al. 2017) (Fig. 1E). We speculated that these motifs may identify binding sites for non-NHR-25 transcription factors to which NHR-25 may tether through protein-protein interactions; such tethering has been described for other nuclear hormone receptors (Nissen and Yamamoto 2000). Alternatively, the motifs may indicate other factors required at the same regulatory region for combinatorial transcriptional action. To this end we performed motif comparison with known databases (JASPAR and cisBP) (Weirauch et al. 2014; Rauluseviciute et al. 2024). Five of the six de Novo motifs had a highly significant match to known *C. elegans* transcription factor motifs when FDR was taken into account (Gupta et al. 2007) (Fig. 1E). For example, a GAGA motif was enriched which significantly matched the known EOR-1 binding motif (p = 4.87e-06) – a protein also expressed in vulval cells and non-seam hypodermal cells (D. Thurtle-Schmidt unpublished observations; Howard and Sundaram 2002).

Previous studies performed ChIP-seq on a tagged version of NHR-25 derived from a low-copy fosmid integrant at L1 (Shao et al. 2013) and L2 (Araya et al. 2014), thus we sought to compare our enriched peaks to these studies. For both datasets we called peaks from the alignments using the same stringent process as outlined above. For the L2 dataset from Araya et al, we identified a greater total number of peaks, 7693 after the filtering process, which is similar to the number of peaks identified for this dataset in a different study (Katsanos and Barkoulas 2022). In comparison we observed significant overlap between our observed enriched peaks to this L2 set from Araya et al: 900 peaks were shared (Jaccard index = .0809, Monte Carlo Simulation p < .0009) (Supplementary Fig. S1C). This concordance in peaks suggests consistent binding between L1 and L2 stages for NHR-25. In a similar analysis performed for the L1 ChIP-Seq dataset from Shao et al, we identified 1173 total peaks with 111 peaks overlapping with our peak set (Jaccard index = .0261) (Supplementary Fig. S1D). While this percentage was still a significant overlap (Monte Carlo Simulation p < .0009), it was less overlap than the L2 dataset. However, the original publication (Shao et al. 2013) did not recover the known NR5A2 binding motif from their peak set suggesting possible non-specific or aberrant enrichment was captured.

### NHR-25 shows distinct distal and proximal binding profiles

Eukaryotic transcription factors appear to bind and act either proximal to the promoter, or distally, from sites far from the promoter (Ray-Jones and Spivakov 2021). In *C. elegans*, the role for more distal regulatory elements has been underappreciated due to the highly compact nature of the *C. elegans* genome resulting in short intergenic sequences and regulatory focus on regions directly upstream of promoters (Blumenthal and Spieth 1996; Reinke et al. 2013). More recent genomic explorations have shed light on distal regulatory elements in *C. elegans* (Araya et al. 2014; Gerstein et al. 2010; Evans et al. 2016; Daugherty et al. 2017; Jänes et al. 2018). Thus, we investigated if there was evidence for both proximal and distal binding and action for a specific transcription factor, NHR-25. In *C. elegans*, chromatin has relatively stable domains across larval development which can be categorized into twenty different chromatin states across the genome (Evans et al. 2016). Thus, for the 1140 enriched NHR-25 regions identified, we investigated which chromatin types were most enriched in the regions as compared to the background genome distribution (Figure 2A). We found that four chromatin types were enriched among the NHR-25 peaks: types 1 and 2, which includes promoter (type 1) and 5’ proximal and gene body (type 2) classified chromatin, as well as type 8 and 9, which are intronic enhancer (type 8) and intergenic enhancers (type 9). All other chromatin types were depleted or not significantly enriched or depleted relative to genomic background. This analysis suggests that NHR-25 may exhibit two binding modes – promoter binding and enhancer-type binding. To determine if the enhancer-type binding is indicative of more action-at-a distance function we identified the closest transcription start site to each peak and compared the average absolute distance based on the two chromatin types. Those peaks that were in promoter-type NHR-25 enriched regions (types 1 and 2) were closer to transcription start sites than the enhancer-type (types 8 and 9) NHR-25 enriched regions (one-sided T-test, t = -8.6095, p < 2.2e16) suggesting NHR-25 may have two binding or functional modes: promoter proximal and distal regulatory elements (Fig. 2B).

**Figure 2.**
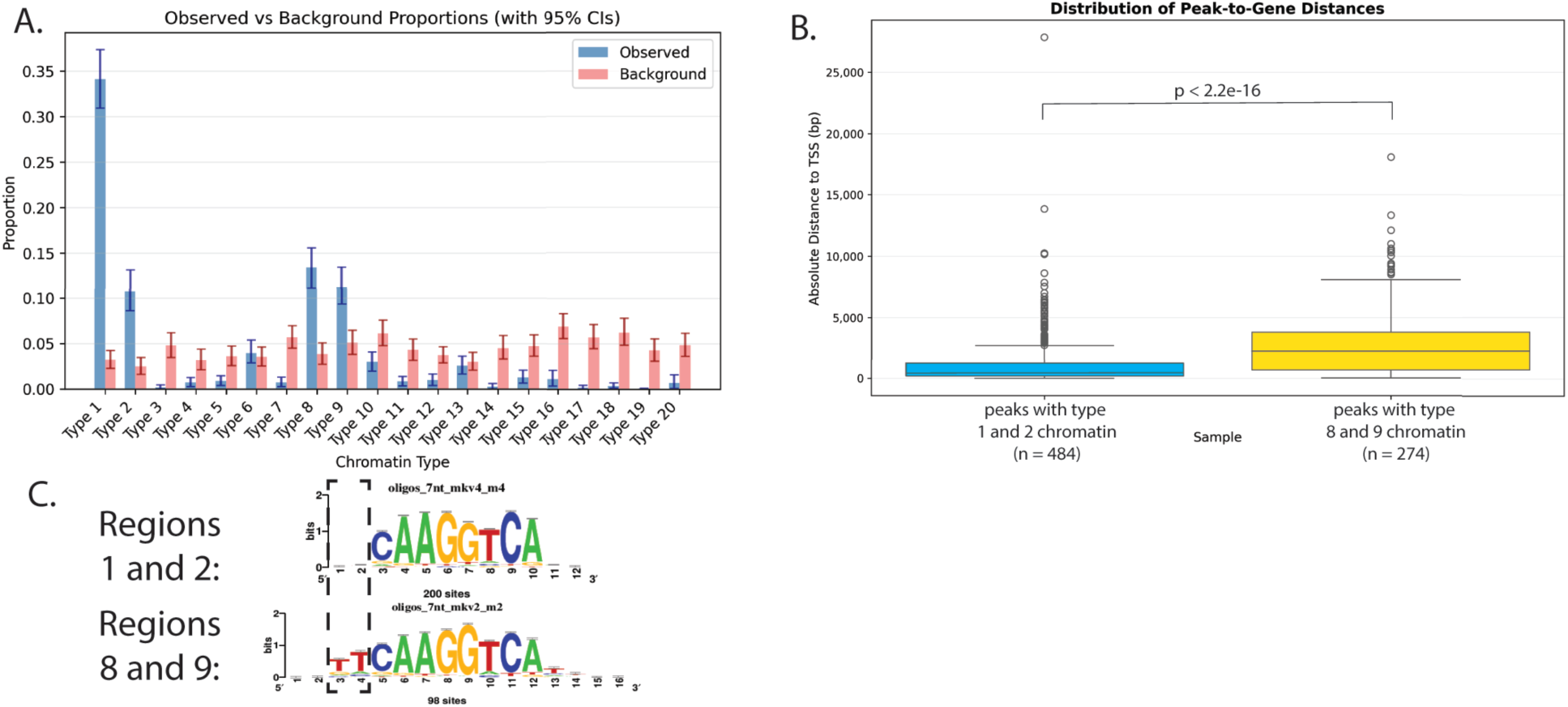
NHR-25 binds two distinct chromatin contexts. (A) Bootstrap analysis of NHR-25 identified binding regions as compared to the 20 chromatin types identified in Evans et al. Error bars represent 95% confidence intervals. (B) Distance from the middle of each binding peak identified to the nearest transcription start site separated by chromatin type. P-value represents 1-sided t-test. (C) RSAT motif analysis of the NHR-25 peaks separated by chromatin type. The dashed box shows the extended bases in the motif identified from peaks with either 8 or 9 type chromatin as defined in Evans et al.

As the underlying chromatin reveals two different binding modes for NHR-25 we also sought to determine if there was also an underlying sequence difference indicative of these different binding actions and navigation of the chromatin environment. To identify sequence characteristics of proximal vs distal NHR-25 enriched regions, we split those peaks that were in one of two classified chromatin promoter (1-2) or distal (8-9) chromatin regions and performed de novo motif discovery and enrichment on each group of peaks (Nguyen et al. 2018). NHR-25 motifs were identified from each group of peaks; however, the motif recovered from enhancer-type chromatin (region 8 and 9) was longer with the full TCA extension present compared to the promoter-type NHR-25 motif identified (Fig. 2C). Since de novo motif identification can be influenced by motif algorithm, we sought to confirm this difference by performing the same analysis with a different motif algorithm (Grant and Bailey 2021). Similarly, the regions from enhancer-type chromatin (regions 8 and 9) recovered the entire NHR-25 binding motif (supplementary Fig. 2), whereas the de novo motif discovered for promoter-type chromatin consisted of only the central 6 bp. Thus, both motif identification programs identified motifs with extended proximal extensions in the enhancer-type chromatin suggesting different sequence recognition by NHR-25 of its DNA binding motif depending on the chromatin context. The de novo motif analysis also identified additional motifs for each chromatin context not likely bound by NHR-25. As analyzed through matrix clustering, some of these motifs are unique to the different types of chromatin, perhaps implying chromatin-context specific differences in transcription factors to which NHR-25 associates via protein-protein interactions (Supplementary Fig. S2B) (Castro-Mondragon et al. 2017).

### NHR-25 regulates metabolic and stress genes

*nhr-25* is an important developmental regulator critical for epidermal specification, vulval development, and molting (Gissendanner and Sluder 2000; Chen et al. 2004; Silhánková et al. 2005). In the larval stages it is expressed in hypodermis, seam cells, vulva, and a subset of neurons (Gissendanner et al. 2004; Silhánková et al. 2005; Cao et al. 2017). Thus, we sought to determine the genes *nhr-25* regulates at a given time in development, to elucidate the molecular underpinnings of the well-characterized mutant phenotypes. As *nhr-25* is an essential gene, we used a conditional knockdown and a hypomorphic mutant to characterize *nhr-25*-dependent transcriptional changes (Gissendanner and Sluder 2000; Chen et al. 2004). Specifically, we measured the transcriptional profile through an *nhr-25* mutant with a single amino acid change in its DNA binding domain, which reduces NHR-25 binding to the canonical NR5A2 motif, and knockdown of *nhr-25* through RNAi (Fire et al. 1998; Chen et al. 2004). N2 and *nhr-25(ku217)* worms were grown and synchronized at the L1 stage and plated on food: N2 worms were plated on L4440 control RNAi bacteria and bacteria targeting *nhr-25* for knockdown. For direct comparison, *nhr-25(ku217)* worms were plated on L4440 control bacteria. To identify substantial changes following gene knockdown by RNAi which was introduced at L1, worms were harvested at L3 and prepared for sequencing in biological triplicate. Since *nhr-25* is a heterochronic gene and can result in developmental delays (Hada et al. 2010), we wanted to ensure that worms were harvested at the same developmental time point for each genotype. To that end, we picked 10 worms per genotype and replicate for each RNA-seq sample ensuring comparable developmental stage by visual inspection, paying special attention to the gonad and vulval morphology.

To determine the *nhr-25* transcriptome, we performed differential expression analysis between each mutant, the point mutation and RNAi knockdown, and wild type. For *nhr-25(RNAi)*, 752 genes were differentially expressed with 354 showing upregulation and 398 genes showed downregulation (p-adj < .05, Fig. 3A, Table S2). Similarly, in the *nhr-25(ku217)* point mutant 1787 genes were differentially expressed of which 1156 were upregulated and 631 genes were downregulated (p-adj < .05, Fig. 3B, Table S2). There was high concordance between datasets with 265 genes differentially expressed in both conditions (Fig. 3C) (Fisher’s exact test p < .05). These observed widespread transcriptional changes demonstrate NHR-25’s role as a key cell-fate transcriptional regulator. To determine which molecular processes are affected by *nhr-25* we performed GO term analysis using wormCAT for all upregulated genes (Fig. 3D) and all downregulated genes (Fig. 3E) across both conditions (Higgins et al. 2021). Genes involved in metabolism and stress response were enriched in both the downregulated and upregulated differentially expressed genes. This finding is consistent with previous microarray studies of *nhr-25(RNAi)* (Ward et al. 2014a). Additionally, transcription factors were specifically enriched in downregulated genes suggesting that NHR-25 normally upregulates a transcriptional cascade. Major sperm protein genes were enriched in the upregulated gene set suggesting a possible role for *nhr-25* normally repressing this transcriptional program prior to the onset of spermatogenesis.

**Figure 3.**
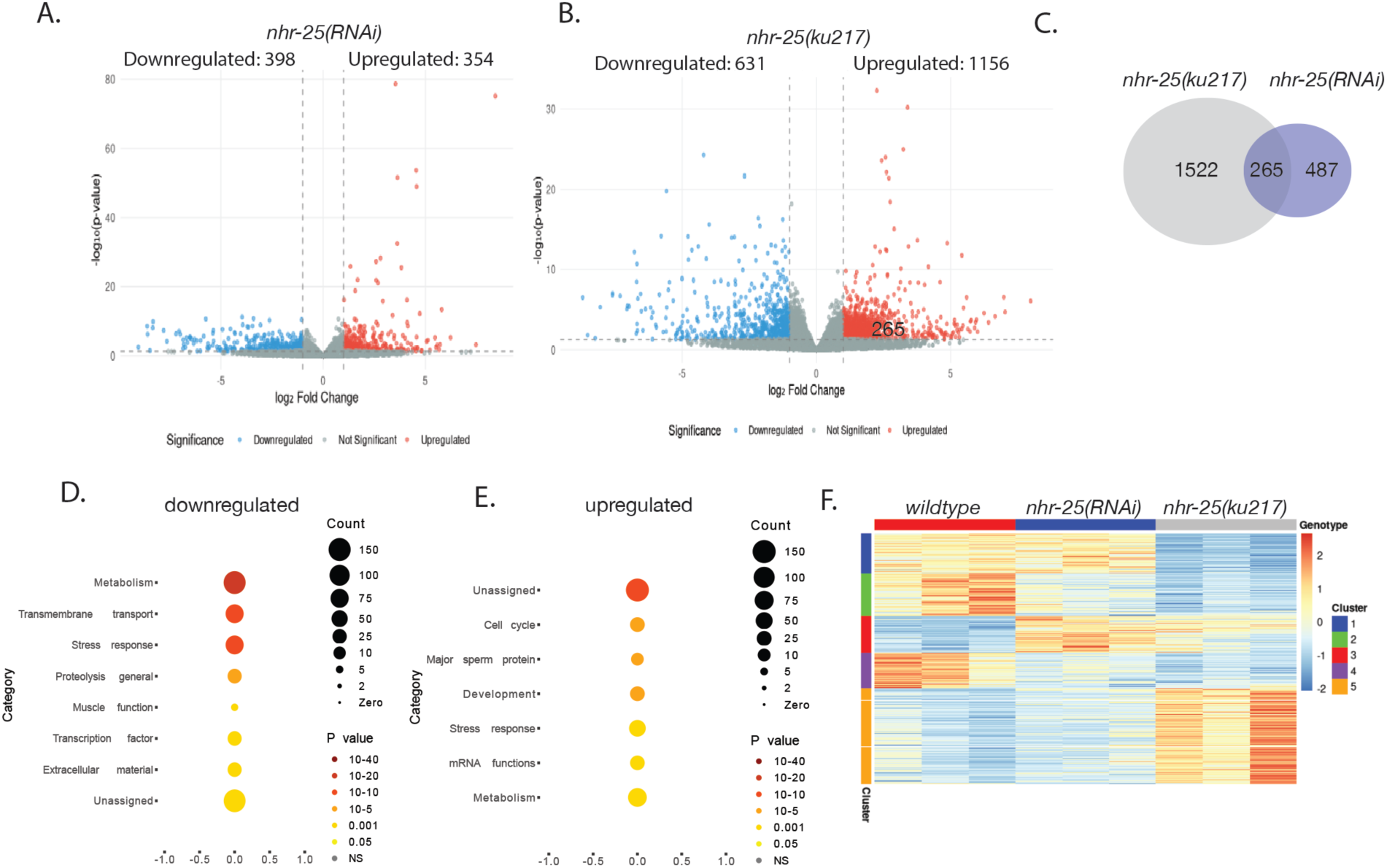
Differential expression analysis of *nhr-25* point mutant and *nhr-25(RNAi)*. (A and B) Differentially expressed genes of *nhr-25(RNAi)* compared to wild type (A) and *nhr-25(ku217)* compared to wild type (B) at L3. (C) the number of shared and unique differentially expressed genes between the two conditions. (D and E) WormCat analysis of the upregulated (C) and downregulated (D) differentially expressed genes across both conditions (Higgins et al. 2021). (E) Z-score normalized heatmap of all differentially expressed genes identified across both conditions in each sample. Cluster analysis of all differentially expressed in either the *nhr-25(ku217)* point mutant or *nhr-25(RNAi)* genes across all three conditions and clusters defined and ordered through K-means clustering analysis with K = 5.

The *nhr-25(ku217)* point mutant was originally isolated in a screen for defects in epidermal development (Chen et al. 2004). It is a single amino acid mutation (L32F) in the DNA binding domain of *nhr-25* and *in vitro* reduces binding of NHR-25 to the canonical NR5A2 motif. As *nhr-25 (ku217)* is a hypomorphic mutant allele, we hypothesized that the regulation of target gene transcription may be similar to *nhr-25* RNAi knockdown. To determine if the mutants showed distinct or more similar expression profiles, we performed principal component analysis. Although there was some variability in the wild-type biological replicates, the PCA showed clear separation between the three genotypes, especially on PC2 (supplemental Fig. S3A). This result indicated that although both RNAi and the point mutant are hypothesized to reduce *nhr-25* function the transcriptional outcomes are at least somewhat distinct.

Since the PCA analysis showed distinct transcriptional profiles for each condition, we were interested in identifying the major expression clusters between *nhr-25(RNAi)* and *nhr-25(ku217)*. To determine the optimal number of clusters, we applied the gap statistic method, which indicated that k = 5 provided the most robust clustering of the data (Supplemental Fig. S3B) (Tibshirani et al. 2001). Visualization of the clusters revealed that genes in clusters 1 and 5 showed distinct expression patterns in the *nhr-25(ku217)* point mutant compared to both *nhr-25(RNAi)* and wild type (Fig. 3F, Table S3). Specifically, the genes in cluster 1 had very low expression in *nhr-25(ku217)* with higher expression in both wild type and *nhr-25(RNAi)*. Cluster 5 showed the opposite trend in expression: high expression in *nhr-25(ku217)* and lower expression in both wild type and *nhr-25(RNAi)*. These distinct expression groups suggest that the point mutant may not function as a simple hypomorph to reduce *nhr-25’s* function.

To identify if there were shared functions in the gene clusters and possible candidate target genes specific to the *nhr-25(ku217)* mutant we performed WormCat enrichment analysis on the clusters that showed differential expression specific to the mutant (Higgins et al. 2021). Enrichment analysis of the genes in cluster 1 showed enrichment for other transcription factors, specifically nuclear hormone receptors, in addition to lipid metabolism and genes involved in stress response (Fig S3C and Table S3). In contrast, the genes in cluster 5, which showed upregulation specific to *nhr-25(ku217)*, were enriched for genes involved in somatic development and unassigned genes in the *C. elegans* genome (Fig S3D and Table S3). Thus, this analysis showed distinct targets which may be differentially sensitive to impaired NHR-25 function, specifically binding and motif recognition or indicating gain-of-function and new regulatory targets for the mutant.

### Identification of putative direct NHR-25 target genes

This transcriptomic analysis revealed the molecular processes regulated by *nhr-25*. However, RNA-seq shows both direct and indirect targets of *nhr-25*, especially with steady-state transcriptional snapshots as described here. Thus, we were interested in correlating NHR-25 enrichment sites to genes that change expression to identify putative direct targets of NHR-25. For each of the 1140 enrichment peaks identified for NHR-25, we correlated each peak with the closest transcription start site (O’Connor et al. 2020). Next, we compared the closest assigned genes with the differentially expressed genes in either the mutant or RNAi knockdown and identified 107 candidate genes potentially directly regulated by NHR-25 (Table S4). From these potential direct NHR-25 targets, we determined if there was evidence for NHR-25 as an activator, repressor, or dual action transcription factor. Of the 107 identified targets, roughly equal number were activated and repressed by NHR-25, suggesting that NHR-25 is a dual-action transcription factor (Fig. 4A). Since we had identified the two differential binding modes for NHR-25 – promoter proximal and distal – we sought to determine if the activated or repressed genes showed correlation for one type of chromatin as compared to the other. For genes putatively activated by NHR-25, both distal and promoter proximal chromatin type associated in roughly similar proportions. The repressed chromatin was associated with promoter proximal chromatin more than distal (Fisher’s Exact Test, p = .01482), suggesting possible contribution of chromatin context to activator or repressor function.

**Figure 4.**
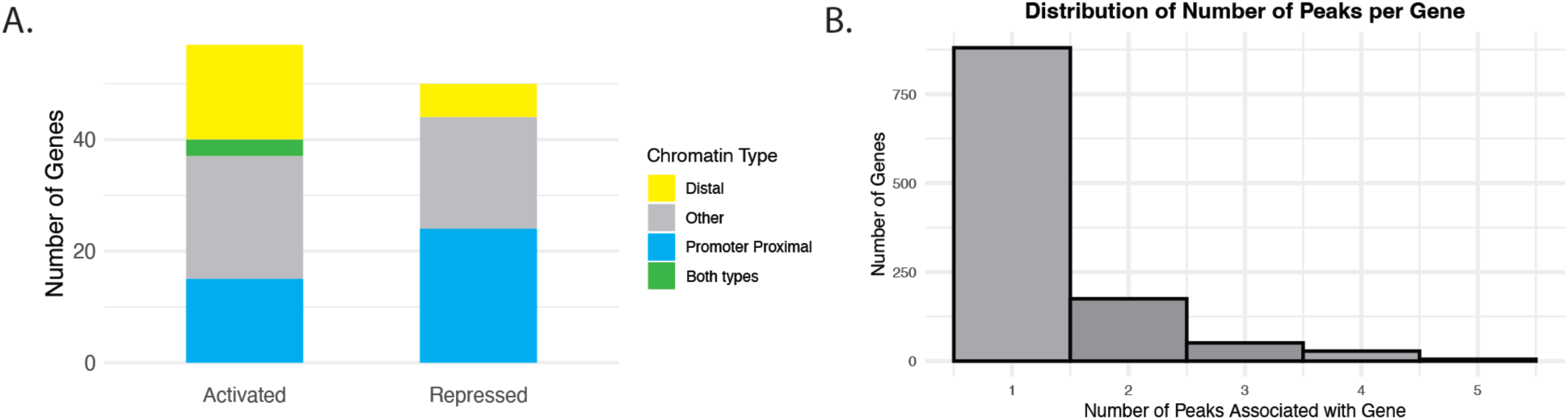
NHR-25 is associated with transcriptional activation and repression. (A) The closest transcription start site was identified for every each NHR-25 enrichment peak and compared to the differentially regulated genes to identify putative direct targets of NHR-25 action. Those genes were split into those that were downregulated in the RNA-seq (activated normally by *nhr-25*) and those that were upregulated in the RNA-seq (repressed normally by *nhr-25*). (B) Histogram of the number of peaks associated with each identified gene.

A notorious challenge in understanding transcription factor function is correlating where a transcription factor binds to its target gene. A recent study showed low correlation between transcription factor binding and function in yeast, an organism with a similarly compact genome (Mahendrawada et al. 2025). With the potential direct NHR-25 targets we identified, we investigated if there were any features from the ChIP-seq data that could predict direct targets. We noted that the peaks associated with the putative direct NHR-25 targets varied in chromatin type, distance (in base pairs) to the TSS of the target genes, and number of ChIP peaks associated with a single target gene (Fig. 4B). Using a logistic regression model, we found that the number of peaks was a highly significant predictor of differential gene expression (Estimate = .543, SE = .1, p = 5.66e-08). Specifically, for every additional peak associated with a gene, the odds of that gene being differentially expressed increased by approximately 1.72-fold. However, neither the distance of the nearest peak to the TSS nor the type of chromatin (promoter proximal (types 1 and 2) or distal (types 8 and 9)) showed a significant effect or improvement of the null model, suggesting that these factors were not good predictors of direct transcriptional targets.

### A single response element regulates multiple genes

As only a subset of the identified ChIP-seq peaks were directly adjacent to a differentially expressed gene when measured through depletion or mutation of the *trans*-acting factor, NHR-25 itself, we were interested in ascribing function to an identified NHR-25 occupied region by testing the putative cis-regulatory regions directly. We identified an NHR-25 enriched peak of interest with type 9 chromatin (Evans et al. 2016) in a large intergenic region that was highly conserved (Felsenstein and Churchill 1996) (Fig. 5A). Interestingly, this intergenic region had multiple NHR-25 peaks clustered nearby (Fig. 5A), which according to our model above suggested functional binding and direct regulation of a nearby target gene. However, neither the directly adjacent genes, *tag-353* or *blmp-1*, were differentially regulated in either the RNAi knockdown or *nhr-25(ku217)* point mutant transcriptome analyses. This peak contained an NR5A2 motif as well as GAGA motifs (Fig. 5B), both of which were highly enriched in our overall NHR-25 ChIP-seq dataset (Fig. 1E). To test the function of this DNA sequence directly, we used CRISPR/Cas-9 mutagenesis and isolated two distinct, but adjacent alleles through homology-directed repair and non-homologous end joining: an 11-bp deletion (Δ11) near the summit of the identified peak directly adjacent to a putative NHR-25 binding motif and a 49-bp deletion which deleted the NHR-25 motif itself (Δ49) and overlapped a portion of the Δ11 (Fig. 5B). These mutant alleles allow us to directly test the function of a cis-regulatory region identified through ChIP-seq.

**Figure 5.**
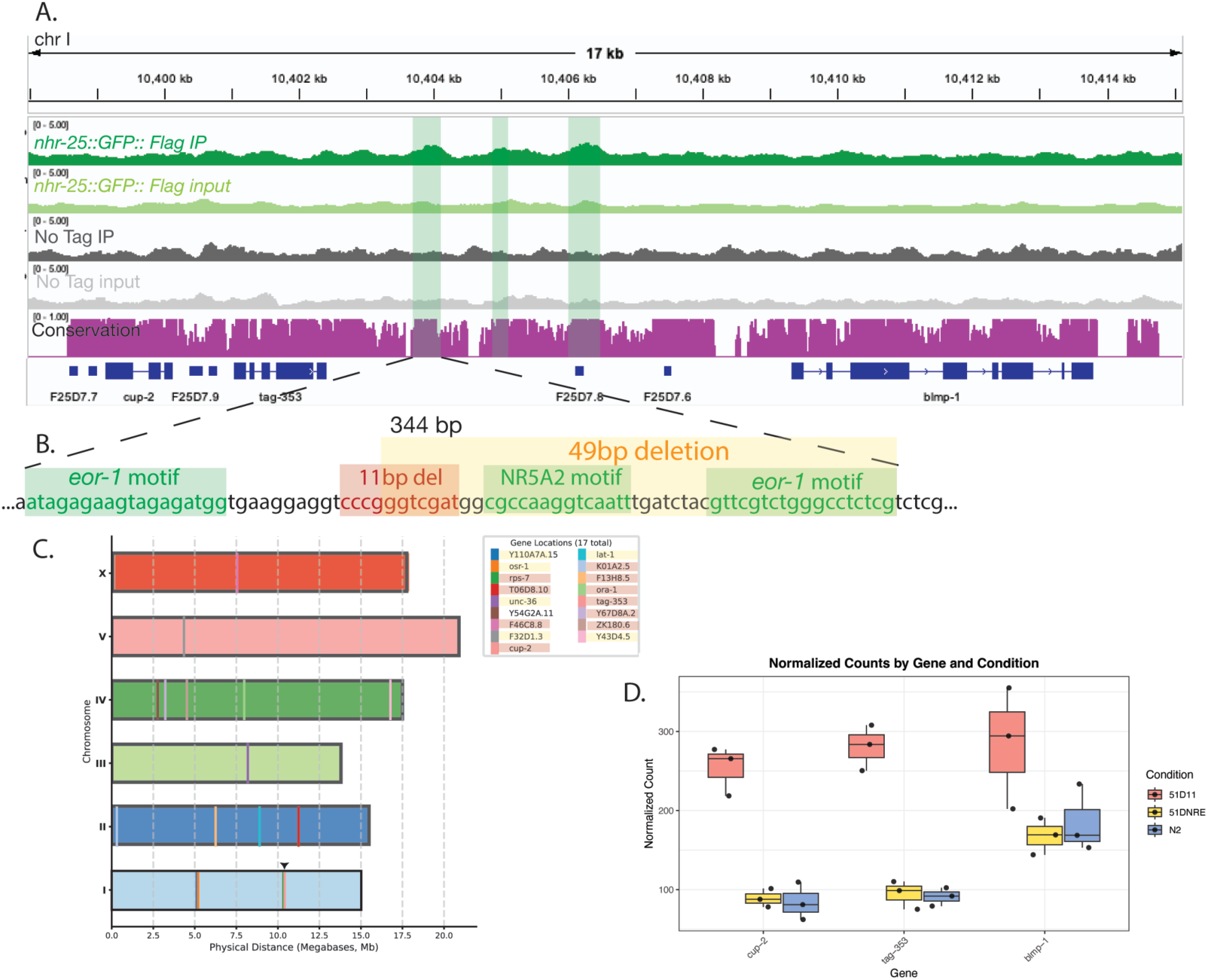
A single response element regulates multiple genes. (A) IGV browser snapshot of *nhr-25:*:*GFP::BioTag::AID*::3xFLAG*, input and no tag IP and input. The bottom magenta track shows conservation across nematodes. The green shaded regions indicated NHR-25 peaks and the peak closest to *tag-353* was mutated. (B) sequence of the NHR-25 enriched region showing the identified motifs and the 11-bp and 49-bp deletions generated with CRISPR-Cas9 mutagenesis. (C) The location of the 18 genes differentially expressed in the Δ11 and Δ49 mutants. Gene names are shaded red if only differentially expressed in the Δ11 mutant and shaded yellow if differentially expressed in the Δ49. One gene is differentially expressed in both mutant conditions and not shaded. The location of the mutations is shown with a black arrowhead above chromosome I. *cup-2 and tag-353* are represented by the same color bar as their position cannot be distinguished at this scale. (D) Normalized counts from RNA-seq of the three adjacent genes in the wild type and 11-bp and 49-bp deletions.

To identify target genes of this cis-regulatory element, we performed RNA-seq on each of these homozygous deletion mutants at L3 and compared the expression to wild-type worms. RNA-seq offers the advantage of not pre-biasing us to the proximal genes and kept the possibility open of long-range transcriptional regulation. Overall, the expression profiles were extremely similar between all the datasets as neither PC1 or PC2 separated the samples by genotype and most of the variance was from differences in the wild-type biological replicates (Supplemental Fig. S4A). This finding was consistent with a mutation in a highly specific, single response element as compared to the broader effects of mutation of the *trans*-acting factor. Through differential expression analysis, 11 genes showed statistically significant differential expression (padj < .01) in the 11-bp deletion whereas 7 genes were differentially expressed (padj < .01) in the 49-bp deletion (Table S5). Of the 18 genes that were differentially expressed in the two mutants only one was shared, *Y54G2A.11*. Surprisingly this gene is not located on Chr I, as expected for mutation of a *cis*-regulatory element, but Chr IV, suggesting possible trans and inter-chromosomal regulation or a secondary effect.

Of the remaining 17 genes that were differentially expressed, we focused our attention on the genes near the cis-regulatory deletions. Two of the genes near the deletions, *cup-2* and *tag-353,* were upregulated in the 11-bp deletion compared to wild type (Fig. 5D, padj < .05). The other adjacent gene, *blmp-1* showed no statistically different expression by RNA-seq, however upon examination of the normalized plot counts *blmp-1* also showed slight upregulation (Fig. 5D). Surprisingly, this upregulation was specific to the 11-bp deletion and not the 49-bp deletion. We conducted qPCR of both mutants and quantified expression of all three of these nearby genes to confirm this finding through a complementary RNA quantification approach with additional biological replicates. As measured by qPCR, *blmp-1* and *tag-353* showed upregulation in the 11-bp deletion allele, but not the 49-bp deletion (Supplemental Fig. S4B), and *cup-2* showed only slight increase in expression over wildtype. Each gene showed high variance in expression as well, possibly capturing stochastic transcriptional consequences of the deletion and a general increase in expression of adjacent genes due to the mutation. Interestingly, *cup-2* but not *tag-353* was also upregulated when *nhr-25* was knocked down by RNAi, linking the *cis*-regulatory deletion and NHR-25. Through this mutational analysis we identified a direct transcriptional role of a highly-specific cis-regulatory region responsible for regulating multiple genes in the native chromatin context.

## DISSCUSION

Here we have described the binding landscape of a highly-conserved, essential nuclear hormone receptor, NHR-25, at endogenous levels in *C. elegans*. Previous studies of NHR-25 genomic binding relied on multi-copy integrants (Araya et al. 2014; Shao et al. 2013) or complementary, but indirect methods (Katsanos and Barkoulas 2022) of identifying protein localization. Our study profiles NHR-25 at endogenous levels revealing two different binding modes for NHR-25 that correlate with chromatin context and promoter proximity. Additionally, through integrated RNA-seq and response element mutagenesis analysis we identified functional consequences of NHR-25 binding and the cis-regulatory element itself, providing mechanistic insights into an essential developmental regulator and mechanisms of transcriptional regulation in *C. elegans*.

Transcription factors appear to function combinatorially. Screens in mammalian systems indicate that few transcription factors display strong activity from regulatory loci containing only their binding motif (Sahu et al. 2022; Alerasool et al. 2022). However, identifying combinatorial partners in transcription factor function is difficult. Through motif analysis we identified non-NHR-25 motifs clustered with NHR-25 motifs in NHR-25-enriched regions, suggesting possible combinatorial partners with NHR-25. In principle, NHR-25 might also tether through protein-protein interactions to a non-NHR-25 transcription factor bound to its motif, or vice versa – a non-NHR-25 transcription factor might tether to a motif-bound NHR-25. However, a previous yeast two hybrid screen failed to identify transcription factor associations with NHR-25, perhaps indicating that combinatorial actions of these factors more likely operate through their binding to clustered motifs within response elements (Ward et al. 2013). These composite response elements may be critical for differential NHR-25 action in distinct cell types as has been proposed for other nuclear receptors (J N Miner and K R Yamamoto 1991).

One potential NHR-25 combinatorial regulator is EOR-1, which binds the enriched GAGA motif. Orthologs of EOR-1 in humans (PLZF) and flies (GAF) have pioneer factor activity that facilitates transcriptional regulation (Judd et al. 2021; Gaskill et al. 2021; Poplineau et al. 2019). Similarly in *C. elegans*, the GAGA motif is enriched at dynamic chromatin accessibility sites in development and the motif is found on longer, nucleosome sized fragments in an ATAC-seq experiment (Daugherty et al. 2017). Additionally, EOR-1 associates with chromatin remodelers in neurons in *C. elegans* (Shinkai et al. 2018). Thus, EOR-1 may be necessary for a subset of NHR-25 transcriptional regulation possibly altering the chromatin landscape to facilitate NHR-25 binding. Notably, the response element edited here contained two GAGA motifs flanking the putative NR5A2 motif illustrating the prevalence and possible synergistic activities of the two factors. Similarly, a previous study identified BLMP-1 as a putative pioneer factor that may specify the amplitude and duration of molting-related gene expression, possibly indicating a general mechanism for regulation of developmental gene expression in *C. elegans* (Stec et al. 2021).

We found that NHR-25 binding sites were enriched for two main classes of chromatin: promoter-type chromatin (types 1 and 2) and enhancer type chromatin (types 8 and 9) (Evans et al. 2016). Through motif analysis of NHR-25 binding in the different types of chromatin, we determined that the NHR-25 binding motif differs in promoter-type chromatin as compared to more distal, regulatory sites. Specifically, the motifs from promoter-type chromatin (1 and 2) lacked the full TCA extension. Our finding of different motifs dependent on chromatin context is consistent with a recent study of the mouse ortholog, NR5A2, which described an NR5A2 motif with a more degenerate TCA extension in open chromatin compared to closed chromatin (Kobayashi et al. 2024). These results suggest that NHR-25 and related orthologs may require different affinities for effective binding or increased kinetic interactions in different chromatin contexts, driving transcription factor function.

We interrogated functional consequences of transcription factor action through two complementary approaches: altering the *trans*-acting factor, *nhr-25* itself, and altering a *cis*-regulatory response element to observe transcriptional consequences. Unsurprisingly, altering the *nhr-25* leads to larger, more global widespread changes in transcriptional activity with enrichment for functions consistent with *nhr-25* mutant phenotypes. Interestingly, comparison of an *nhr-25* DNA Binding Domain (DBD) mutant specific transcriptional changes to the knockdown through RNAi revealed mutant-specific classes of misregulation. Previous studies identified mutations in the DBD of orthologous transcription factors altering their binding in vivo. For example, a naturally occurring variant of SF-1 is found to alter its binding (Ito et al. 2000). Our RNA-seq data suggests this may be the case for *nhr-25(ku217)* mutant as well. Alternatively, the mutant may result in lower transcription factor protein expression due to misfolding and degradation, possibly altering binding kinetics and action. However, these two potential models are not mutually exclusive and suggest either novel binding sites or sites differentially sensitive to NHR-25 binding and function.

Depletion of *nhr-25* did identify putative direct targets including nuclear hormone receptors, potentially revealing important cell-fate gene regulatory networks. One such differentially expressed gene with adjacent NHR-25 binding is *nhr-85*, which is involved in the heterochronic gene network through regulating *lin-4 and* when mutated shows similar egg-laying defects to *nhr-25* mutants, indicating a possible direct transcriptional cascade (Gissendanner et al. 2004; Kinney et al. 2023). Notably, *nhr-85* had multiple NHR-25 ChIP-seq peaks associated with it (Fig. 1D), which our model indicated as the best predictor of a direct transcriptional target. Consistent with this model, other studies in *C. elegans* report target genes of interest with multiple ChIP-seq peaks of specific transcription factors near those target genes (Kinney et al. 2023; Ragle et al. 2025). While this study identified putative direct targets of NHR-25, this model likely underestimated identification of NHR-25 direct targets. First, the model was dependent on the assumption of correlating the transcription factor to the closest transcriptional start site. However, this assumption is likely limiting as we demonstrated through mutation of the regulatory region itself that NHR-25 appears to regulate transcription of remote genes (Fig. 5). Additionally, RNA-seq quantified steady-state expression at a single time point, likely underestimating NHR-25 regulation of dynamic gene expression dependent on the time-point assayed. Despite these limitations, we identified one strong predictor of direct transcriptional targets for NHR-25 that can likely be applied across transcription factors.

This analysis not only interrogated the NHR-25 transcription factor but also a cognate response element itself within its native chromatin context to ascertain the contribution of DNA-sequence directly to transcriptional regulation. ChIP-seq resolution is determined by shearing size, which usually results in identified peaks around 200 – 500 bp, which is much larger than functional response elements, even when considering composite response elements. Previous studies of cis-regulatory deletions in *C. elegans* deleted the entire ChIP-seq peaks to measure specific phenotypic consequences of transcription factor binding (Kinney et al. 2023; Ragle et al. 2025). However, in this study we generated highly-specific, short deletions at the summit of an identified ChIP-seq peak and measured the deletion’s effects through highly sensitive RNA-seq which identified any transcriptional consequences of peak summits directly. For both deletion alleles, RNA-seq revealed a small subset of differentially regulated genes across the chromosomes, suggesting possible *trans*-chromosomal consequences of deleting a non-coding regulatory element. Many of the genes differentially expressed are in the central regions of the chromosomes, consistent with the active, central regions of chromosomes interacting preferentially (Bian et al. 2020). However, the distal chromosomal arms are gene poor in *C. elegans* and thus by chance less likely to harbor any gene, let alone differentially expressed genes. Only one gene was differentially expressed in both deletions. Since the deletions partially overlap, this finding suggests that the regulatory consequences are highly specific within a composite response element. Additionally, identifying transcription factor binding and association through methods with higher resolution such as CUT&RUN and similar methods could also isolate specific response elements within the larger ChIP-seq peaks and direct further studies of specific cis-regulatory mutations (Skene and Henikoff 2017; Kaya-Okur et al. 2019; Emerson and Lee 2022).

In analyzing the transcriptional consequences of the response element deletions, we also examined those genes adjacent and in cis with the generated deletions. Surprisingly, the transcriptional consequences of the adjacent genes were specific to the 11-bp deletion and not to the larger mutant (Δ49) that included deletion of the identified NR5A2 and GAGA motifs. The mutated peak summit was near two other NHR-25 identified peaks within this intergenic region, thus the other peaks may act redundantly under the assayed condition and may compensate for loss of NHR-25 direct binding in the 49-bp deletion. However, we focused on transcriptional consequences of the DNA sequence itself and never directly tested if NHR-25 is no longer enriched in either the 11-bp or 49-bp deletions. Thus, NHR-25 or other critical transcription factors such as EOR-1, may still associate at that site in the 49-bp deletion. Future analysis could directly test cis-regulatory mutations to differential binding by NHR-25. The 11-bp deletion did result in upregulation of three nearby genes, suggesting that this specific DNA sequence acts as a general repressor on nearby promoters and deletion of the 11-bp may disrupt binding of an unknown transcription factor. The regulated promoters of the nearby genes in cis (*cup-2, tag-353, blmp-1*) ranged from 3,000 bp – 5,000 bp from the deletions, which demonstrates that a single, specific response element can regulate multiple promoters at a distance.

## METHODS

### *C. elegans* cultures and strains

*C. elegans* were cultured at 25°C using standard protocols unless otherwise noted. The wild type strain used is the N2 Bristol strain (Brenner 1974). The following mutant and transgenic strains were generated in this study: JDW4 (*ieSi57 [Peft-3::TIR1::mRuby::unc-54 3’ UTR, cb-unc-119(+)] II; nhr-25(wrd2[GFP^BioTag-degron-3xFlaG]) X),* DTW14 (*Δ49 I*), DTW17 (*Δ11 I*). All strains are available upon request.

### Microscopy and ChIP-seq sample and library preparation

N2 (no tag) and *nhr-25:*:*GFP::BioTag::AID*::3xFLAG C. elegans* were synchronized using standard sodium hypochlorite protocols and hatched overnight in 50 mls of S-basal+gelatin with shaking at 20°C. In the morning, worms were diluted to 250 mls total of S-basal+gelatin and 6 mls of concentrated HB101. Worms were grown at 25°C with shaking for 6 hours. Prior to harvesting for ChIP, *C. elegans* were anesthetized with 10 mM sodium azide and mounted onto 2% agarose pads and imaged using Zeiss Axioplan microscope. ChIP was performed as previously described (Askjaer et al. 2014) with the following modifications. Briefly, worms were washed in M9+gelatin and frozen in liquid nitrogen in PBS + Protease inhibitors. Frozen worm pellets were ground under liquid nitrogen then worms were fixed in 1.1% formaldehyde in PBS + protease inhibitors for 20 minutes at room temperature. Formaldehyde was quenched with 125mM glycine and samples sonicated using the Diagenode Bioruptor Pico at 4°C on high for 15 min on high, 30 seconds on and 30 seconds off then repeat a second time. Samples were immunoprecipitated with anti-FLAG M2 Magnetic beads (M8823 Millipore Sigma) and incubated overnight at 4°C. Crosslinks were reversed with Proteinase K and treated with RNAase A. Chromatin was purified using the Zymo ChIP DNA clean and concentrator kit (D5205) with two subsequent 25µl elutions. Purified DNA from immunoprecipitates and inputs were prepared into libraries using the BioO qRNA-seq library prep kit adapted for DNA. Paired-end 50bp sequencing was performed on Illumina Hi-Seq 2500.

### ChIP-seq bioinformatic analysis

Reads were quality checked with FastQC (Andrews 2010). The adaptor and first 12bp from the 5’ end of all reads were trimmed from all reads using Cutadapt due to initial low quality (Martin 2011). Reads were mapped to the Ce10 genome using BWA-MEM with default settings (Li 2013) and duplicates marked with Picard (Institute). MACS2 was used to call peaks for each IP and input pair, including the wild type (no-tag) datasets. Peaks were then visually inspected with IGV to identify those in the *nhr-25::GFP::Flag* dataset that overlapped with peaks detected in the no-tag dataset, and those peaks were discarded (Thorvaldsdóttir et al. 2013). Peaks from both NHR-25 IP dataset replicates were combined. Using bedtools intersect those peaks that overlapped with regions designated as XHOT in all contexts (maphot_ce_selection_reg_cx_simP01_all.bed) were removed (Araya et al. 2014; Quinlan and Hall 2010). These refined peaks were used in RSAT peaks-motifs to detect enriched motifs (Nguyen et al. 2018; Thomas-Chollier et al. 2012), using the following parameters: peak sequences were cut to +/- 500 bp on each side, all four motif discovery algorithms were used. Identified motifs were compared to the JASPAR non-redundant core and the cis-BP *C. elegans* databases (version 2) (Castro-Mondragon et al. 2022; Weirauch et al. 2014). Motifs were clustered using RSAT matrix clustering (Castro-Mondragon et al. 2017). Custom python scripts were used to determine chromatin type and enrichment for identified peaks.

### RNA Interference

Feeding RNAi was performed as described (Timmons et al. 2001) with following alterations. dsRNA was induced in liquid culture using 0.4 mM IPTG. Bacteria were then concentrated and seeded on plates also containing 0.4 mM IPTG. Bacteria carrying pPD129.36 without an insert was used for a control. Synchronized L1 larvae were plated on the plates and allowed to grow as indicated below.

### RNA-seq sample preparation

Worms were synchronized using standard sodium hypochlorite methods as previously described. Eggs were hatched overnight at 25°C in the absence of food. Synchronized L1s were moved to food and allowed to grow at 25°C for 24 hours to L3 stage on 6cm plates seeded with appropriate bacteria. At L3 stage, 10 worms per genotype were picked into 40 ul M9 media with gelatin and 360 ul Trizol added. Samples were flash frozen in liquid nitrogen and subsequently subject to four rounds of freeze-thaw cycles in liquid nitrogen and at 37°C. Following addition of phenol, samples were purified using Qiagen RNeasy protocol. Libraries were prepared from the purified total RNA using BioO qRNA-seq library prep kit, using 50bp SE sequencing.

### RNA-seq analysis

Quality of Reads were checked with FastQC and results complied with MultiQC (Andrews 2010; Ewels et al. 2016). Reads were trimmed for poor quality with trimmomatic and access adaptor cut with CutAdapt (Bolger et al. 2014; Martin 2011). Reads were mapped to the Ce11 genome using HISAT2 to check mapping quality (Kim et al. 2015). Trimmed Fastq files were quantified directly with salmon quant. Differential expression was determined using DeSeq2 and analysis in R studio (Love et al. 2014; R Core Team 2022).

### CRISPR methods

The *nhr-25::GFP::BioTag::3xFlag* endogenous tag was generated using the self-excising cassette (SEC) method (Dickinson et al. 2015). 552 bp (5’) and 534 bp (3’) homology arms with 30 bp of backbone homology were PCR amplified from genomic DNA and Gibson cloned into AvrII+SpeI digested pJW1592 (Ashley et al. 2021) to make pJW1597. We generated a Cas9+sgRNA (F+E) vector (pJW1308) targeting the 3’ end of *nhr-25* using Q5 site-directed mutagenesis (NEB) of pJW1219 as described (Dickinson et al. 2013; Ward 2015). We then injected the pJW1597 repair template at 10 ng/µl and the pJW1308 Cas9+sgRNA vector at 50 ng/µl into CA1200 and recovered knock-ins and excised the SEC as previously described (Dickinson et al. 2015). Plasmid and oligo sequences available upon request. pJW1592 is available from AddGene.

The *tag-353 blmp-1* intergenic region was edited using CRISPR-Cas9 system according to (Paix et al. 2014). Briefly, CRISPR-Cas9 ribonucleoprotein (RNP) complexes were preassembled by mixing 110 ng/μl tracr RNA (IDT), 100 ng/μl of each guide RNAs (Alt-R crRNA, IDT) and 250 ng/μl Cas9-NLS purified protein (MacroLab, QB3 Berkeley) and incubating at 37 °C for 10 min. 110 ng/μl of the ssODN donor template (IDT) (Table S6) and 50 ng/μL pRF4: co-injection marker *rol-6 (su1006)* expression plasmid (Kramer et al. 1990) were then added to the complex mixture. The mix was microinjected into the gonadal arms of HL178 worms (Mello and Fire 1995). Injected P0 hermaphrodites were singled and allowed to lay eggs. F1 rollers (Rols) from jackpot-brood plate(s) based on the number of Rol animals in a plate were singled. On the second day after the initiation of egg laying, F1s were frozen individually in worm lysis buffer (10 mM Tris-HCl, pH 8.2, 50 mM NaCl, 2.5 mM MgCl_2_, 0.45% NP-40, 0.05% polyethylene glycol 8000) with 1mg/mL Proteinase K (Ambion) and subjected to single-worm PCR for genotyping. Mutations were screened by MiSeq next generation sequencing (Illumina). Progeny from the positive F1 with desired mutations was recovered and singled to get the homozygous mutant. Isolated mutants were backcrossed with N2 wild-type males two times.

### qPCR sample preparation and analysis

Quantitative PCR was performed on cDNA samples, created from isolated L3 RNA using LunaScript RT Supermix Kit (NEB). RNA was harvested as described for RNA-seq above. The cDNA was then quantified using an Mx3000 qPCR machine (Agilent), with Luna Universal qPCR Master Mix (NEB). For each sample, in addition to the genes tested, three normalization genes were also assayed. To calculate relative gene expression, each test was normalized to the geometric mean of three reference genes (Table S6) as described in (Vandesompele et al. 2002). The ΔΔCt or Livak method was used for normalization to both the internal reference genes and wild-type sample (Schmittgen and Livak 2008).

## DATA ACCESS

The ChIP-seq and RNA-seq data generated in this study have been submitted to the NCBI BioProject database (https://www.ncbi.nlm.nih.gov/bioproject/) under accession number PRJNA1380758.

## COMPETEING INTERESTS

The authors declare no competing interests.

## ACKNOWLEDGEMENTS

We thank members of the Yamamoto lab and the Thurtle-Schmidt lab for insightful discussions. Research reported in this publication was supported by the National Institute of General Medical Sciences of the National Institutes of Health under Award Number R15GM147889. The content is solely the responsibility of the authors and does not necessarily represent the official views of the National Institutes of Health. Additional support was provided by National Science Foundation grant (MCB-1615826). Additionally, DTS was supported by a Gordon and Betty Moore Foundation-sponsored Life Sciences Research Foundation fellowship. Some *C. elegans* strains were provided by the CGC, which is funded by NIH Office of Research Infrastructure Programs (P40 OD010440). JDW was supported by National Institute of General Medical Sciences of the National Institutes of Health (K99GM107345, R00GM107345, and R01 GM138701). *Author contributions:* D.M.T.S and K.R.Y. planned and conceived the study. D.M.T-S and J.D.W. performed ChIP-seq. J.D.W and M.A. generated transgenic lines. D.T.S and A.L.S. performed RNA-seq. D.T.S designed the analysis pipeline. D.T.S and A.L.S analyzed and interpreted the data and wrote the manuscript. All authors discussed the results and commented on the manuscript.

